# High-resolution spatiotemporal transcriptomic maps of developing *Drosophila* embryos and larvae

**DOI:** 10.1101/2021.10.21.465301

**Authors:** Mingyue Wang, Qinan Hu, Tianhang Lv, Yuhang Wang, Qing Lan, Zhencheng Tu, Rong Xiang, Yanrong Wei, Kai Han, Yanru An, Mengnan Cheng, Jiangshan Xu, Miguel A. Esteban, Haorong Lu, Wangsheng Li, Shaofang Zhang, Ao Chen, Wei Chen, Yuxiang Li, Xiaoshan Wang, Xun Xu, Yuhui Hu, Longqi Liu

**Affiliations:** BGI-Shenzhen, Shenzhen 518083, China; Shenzhen Key Laboratory of Gene Regulation and Systems Biology, School of Life Sciences, Southern University of Science and Technology, Shenzhen 518005, China; Department of Biology, School of Life Sciences, Southern University of Science and Technology, Shenzhen 518005, China; College of Life Sciences, University of Chinese Academy of Sciences, Beijing 100049, China; South China University of Technology, Guangzhou 510006, China; BGI College & Henan Institute of Medical and Pharmaceutical Sciences, Zhengzhou University, Zhengzhou 450000, China; BGI-Qingdao, BGI-Shenzhen, Qingdao, 266555, China; Laboratory of Integrative Biology, Guangzhou Institutes of Biomedicine and Health, Chinese Academy of Sciences, Guangzhou 510530, China; CAS Key Laboratory of Regenerative Biology and Guangdong Provincial Key Laboratory of Stem Cells and Regenerative Medicine, Guangzhou Institutes of Biomedicine and Health, Guangzhou 510530, China; Institute of Stem Cells and Regeneration, Chinese Academy of Sciences, Beijing 100101, China; China National Gene Bank, BGI-Shenzhen, Shenzhen 518120, China; Guangdong Provincial Key Laboratory of Genome Read and Write, BGI-Shenzhen, Shenzhen, 518120, China; Shenzhen Key Laboratory of Single-Cell Omics, Shenzhen 518083, China

## Abstract

*Drosophila* has long been a successful model organism in multiple fields such as genetics and developmental biology. *Drosophila* genome is relatively smaller and less redundant, yet largely conserved with mammals, making it a productive model in studies of embryogenesis, cell signaling, disease mechanisms, etc. Spatial gene expression pattern is critical for understanding of complex signaling pathways and cell-cell interactions, whereas temporal gene expression changes need to be tracked during highly dynamic activities such as tissue development and disease progression. Systematic studies in *Drosophila* as a whole are still impeded by lack of these spatiotemporal transcriptomic information. *Drosophila* embryos and tissues are of relatively small size, limiting the application of current technologies to comprehensively resolve their spatiotemporal gene expression patterns. Here, utilizing SpaTial Enhanced REsolution Omics-sequencing (Stereo-seq), we dissected the spatiotemporal transcriptomic changes of developing *Drosophila* with high resolution and sensitivity. Our data recapitulated the spatial transcriptomes of embryonic and larval development in *Drosophila*. With these data, we identified known and previously undetected subregions in several tissues during development, and revealed known and potential gene regulatory networks of transcription factors within their topographic background. We further demonstrated that Stereo-seq data can be used for 3D reconstruction of *Drosophila* embryo spatial transcriptomes. Our data provides *Drosophila* research community with useful resources of spatiotemporally resolved transcriptomic information across developmental stages.

## INTRODUCTION

For over a century, *Drosophila* has been a productive model organism for developmental biologists and geneticists to study a diverse range of developmental processes, such as embryogenesis, organogenesis, gametogenesis, and aging. Numerous studies in *Drosophila* have led to groundbreaking discoveries, which have greatly impacted the biomedical field.

The transcriptomic profiles of cells and tissues largely determine their functions. In multicellular organisms, various types of cells with diverse transcriptomic profiles together orchestrate the functions of tissues and organs to ensure the smooth execution of biological processes. Traditionally, tissue/cell types were distinguished based on their function, anatomy, morphology, and expression of a few marker genes. As a well-established model organism, *Drosophila* has been intensively studied as to its tissue-specific transcriptomes. Marker gene expression profile in each tissue/cell type are readily available. Several databases have been established to curate the collection of tissue-specific transcriptomic profiles in *Drosophila*, including FlyAtlas1 (Chintapalli et al., 2007), FlyAtlas2 (Leader et al., 2018) and DGET (Graveley et al., 2011).

Recently, the rapid development of single-cell multi-omics technologies enables the mapping of genomic, transcriptomic, epigenomic and proteomic information at single-cell resolution. This has led to multiple studies revealing cell heterogeneity of *Drosophila* tissues by single-cell multi-omics sequencing, such as embryo (Karaiskos et al., 2017), imaginal disc (Ariss et al., 2018; Bageritz et al., 2019; Deng et al., 2019), gut (Guo et al., 2019; Hung et al., 2020), brain (Brunet Avalos et al., 2019; Davie et al., 2018) and gonad (Jevitt et al., 2020; Rust et al., 2020; Witt et al., 2019) (reviewed in Li, 2020). These studies substantially expanded our knowledge of cellular diversity, functional heterogeneity and microenvironmental regulation in *Drosophila* tissues. Comparing to traditional cell type identification methods, single-cell multi-omics sequencing measures many more dimensions of cell states and expression profiles, revealing novel cell types in multiple tissues. The Fly Cell Atlas project (Li et al., 2021), a global collaborative effort, is in progress to build comprehensive cell atlases of *Drosophila* tissues across developmental stages with single-cell multi-omics techniques.

Owing to the complex intercellular communication and coordination within or between tissues in multicellular organisms, our understanding of cellular functions greatly relies on their morphological contexts. The spatial information of cells’ transcriptomes provides a wealth of insights for how cells coordinate to perform their biological functions in cell-cell signaling, metabolism and development. Currently, several databases are available for the spatial expression pattern of developing *Drosophila* embryos and larvae, such as Berkeley *Drosophila* Genome Project (BDGP) gene expression pattern studies (Tomancak et al., 2002) and Fly-FISH (Lécuyer et al., 2007). However, these databases mainly utilize *in situ* hybridization, which has several inherit drawbacks, including inability to detect unknown transcripts and isoforms, difficulty in identifying transcripts with low copy number, and lack of accuracy in quantification of gene expression levels. Moreover, the *Drosophila* spatial transcriptome is yet to be completely resolved. Within the ∼14,000 genes in the *Drosophila* genome, over 40% still lack information in their spatiotemporal expression patterns (Zhou et al., 2019), and the ones with these patterns are also far from comprehensively resolved.

Recent years have witnessed the advances of spatial transcriptomic technologies, which utilize various methods to resolve the spatial patterns of transcriptomes, including computational strategies (Karaiskos et al., 2017), physical segmentation (Junker et al., 2014), local mRNA capture and sequencing (Rodriques et al., 2019) etc. These methods vary in throughput, number of genes measured and spatial resolution, and have been successfully applied to multiple tissues (reviewed in Liao et al., 2021). In an effort to capture the actual spatial transcriptomes *in situ*, some methods utilize surfaces covered with an array of uniquely DNA-barcoded beads. mRNAs from tissues (usually a section) are directly transferred to the surface and captured by the beads carrying oligo dT sequences. The spatial patterns of transcriptomes can thus be resolved after sequencing and mapping. These tools enable untargeted and comprehensive capture of cellular transcriptomic profiles *in situ*, thus can overcome the disadvantages of *in situ* hybridization, make valuable supplement to the current spatial transcriptomics databases and facilitate discoveries of previously undetectable transcriptomic changes. They have been successfully applied to tissues such as mouse brain (Ortiz et al., 2020), human brain (Chen et al., 2020) and human heart (Asp et al., 2019).

Most of the existing DNA-barcoding array methods lack sub-micrometer level resolution or sufficient mRNA capture efficiency, impeding their application in *Drosophila* embryos (∼500 μm × 100 μm in section size). Due to the small sizes of these samples, efforts in resolving their complete spatial transcriptome remained computational (Karaiskos et al., 2017). Our recently developed SpaTial Enhanced REsolution Omics-sequencing (Stereo-seq) technique (Chen et al., 2021) can resolve spatial transcriptomes with nanometer resolution while retaining high sensitivity, providing a powerful tool to obtain spatial transcriptome from small-sized samples like *Drosophila* embryos. In Stereo-seq, patterned DNA nanoballs (DNBs) with randomly barcoded sequences are photolithographically etched on a modified chip ∼500 nm away from each other. This technique allows mRNA capture with high density and sensitivity, enabling spatial transcriptomic analysis at a much higher resolution. As a sequencing-based spatial transcriptomic technique with subcellular resolution, Stereo-seq makes it possible to capture the detailed spatially resolved transcriptomes in small-sized samples such as *Drosophila* embryos and larvae.

At 25 °C, the *Drosophila* life cycle starts with ∼1 day of embryogenesis [can be arbitrarily divided into 17 stages based on morphology (Campos-Ortega and Hartenstein, 1997)], followed by three larval stages which take altogether ∼5 days. Pupal stage follows and lasts for another ∼4 days. Adult flies subsequently eclose from the pupal cases. In this study, we applied Stereo-seq to late-stage embryos and all stages of larvae, generating spatial transcriptomic data across these developmental stages. Computational analysis could correlate bin clusters with anatomical structures. With the Stereo-seq data, we also detected subregions of tissues by refined clustering, and identified active transcription factor regulatory networks in multiple tissues across embryonic and larval development. Lastly, we demonstrated that with Stereo-seq data from all the sections of an embryo, it is possible to 3D reconstruct the spatial transcriptome of an embryo *in silico*. This opens exciting opportunities for *Drosophila* research to be done in an organism-wide, 3D spatially resolved and systematic manner.

## RESULTS

### Application of Stereo-seq to *Drosophila* embryos and larvae

We applied Stereo-seq to the following *Drosophila* samples: late-stage embryos (14-16 h and 16-18 h after egg laying, corresponding to stage 16∼17 of embryogenesis, hereafter termed E14-16 and E16-18 respectively) and all three stages of larvae (hereafter termed L1∼L3) **(Fig.1A)**. In each experiment, a patterned 1cm × 1cm Stereo-seq chip was covered with 10-μm thick cryosection slices from each sample. Stereo-seq was performed as previously described (Chen et al., 2021). Sequencing data were processed and integrated to generate 2D spatial transcriptomes of each sample. On average, Stereo-seq captured more than 1,500 unique transcripts representing over 400 genes per bin (bin 20 × 20 for E14-16, E16-18, L1 and L2, equivalent to ∼10 μm × 10 μm; bin 50 × 50 for L3, equivalent to ∼25 μm × 25 μm) across samples **(Fig.S1A, Table S1)**. Based on Stereo-seq data, 2D expression matrices with defined X-Y coordinates and positional expression profiles of each bin were generated. We then performed unsupervised clustering for each 2D expression matrix **(Fig.S1B)**. Clusters were associated with tissue types according to previously reported marker gene profiles, followed by manual validation and annotation. Clustering results were consistent with anatomical structures of embryos and larvae **(Fig.1B-F, Fig.S2)**.

**Figure 1.**
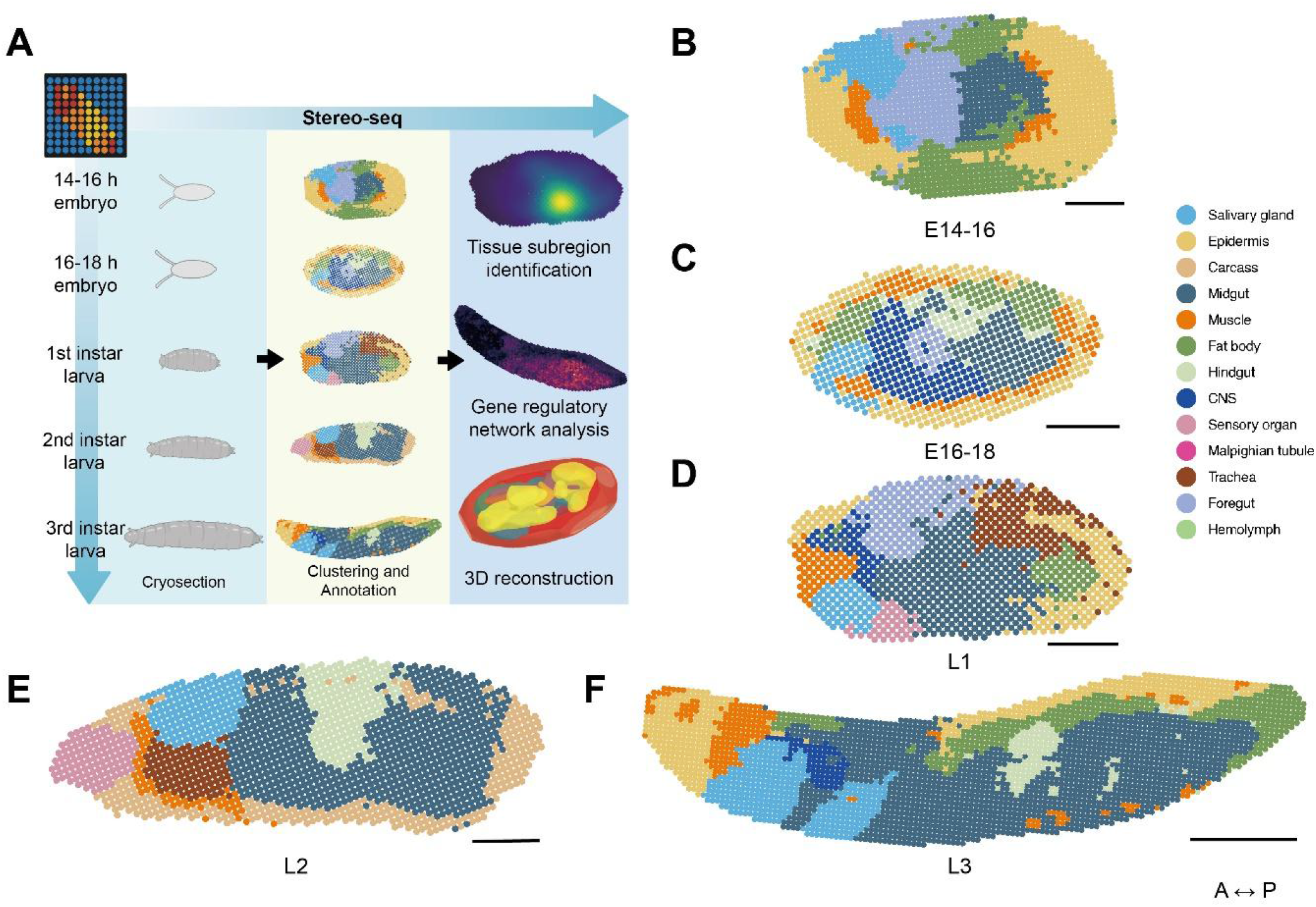
Clustering and annotation of Stereo-seq generated spatial transcriptomes of *Drosophila* late-stage embryos and larvae. **(A)** Schematic workflow of Stereo-seq data acquisition and analysis in this study. **(B-F)** Unsupervised clustering and annotation of representative single sections of **(B)** E14-16, **(C)** E16-18, **(D)** L1, **(E)** L2, and **(F)** L3 samples. Clusters with the same annotation are merged and assigned the same color code. Scale bars = 100 μm for E14-16, E16-18, L1 and L2, and 500 μm for L3. A-P: anterior-posterior.

To validate the spatial transcriptome we obtained with Stereo-seq, we compared the localization patterns of a list of transcripts that we observed in late-stage embryo samples with known *in situ* hybridization results for those transcripts, obtained from BDGP database (Tomancak et al., 2002, 2007). The spatial patterns of these transcripts in our Stereo-seq data matched well with those of *in situ* hybridization results **(Fig.S3 and S4A)**. These observations indicate that Stereo-seq effectively reflects the spatial gene expression patterns of *Drosophila* embryos. In addition, our Stereo-seq data yielded localization information for multiple transcripts that are not included in previous high-throughput *in situ* hybridization experiments from BDGP and Fly-FISH projects **(Fig.S4B)**. These data provide a valuable complement to current *in situ* hybridization databases.

Thus, Stereo-seq generated high-quality spatial transcriptomic data from *Drosophila* embryo and larva samples, identifying spatial patterns of both transcripts matching previous *in situ* hybridization data and previously undetected transcripts.

### Spatiotemporal gene expression profiling reveals subregions of embryonic and larval midgut

By virtue of large number of genes captured by Stereo-seq, clusters corresponding to highly complex tissues can be further subclustered based on differences in cell subtype marker gene expression. The digestive system occupies the largest space in late-stage *Drosophila* embryos and larvae, thus correlated to the largest number of bins in our Stereo-seq data. Moreover, unsupervised clustering generated multiple distinct clusters corresponding to midgut across samples, suggesting tissue heterogeneity within embryonic and larval midgut. Thus, we extracted clusters representing midgut from all stages of investigated samples and performed refined clustering. Further unsupervised clustering of midgut clusters revealed subregions with distinct marker genes **(Fig.S5A)**. We then inspected the expression patterns of marker genes in each subcluster.

In adult *Drosophila* midgut, recent studies characterized the molecular specificity and cellular composition of midgut subregions by bulk and single-cell RNA-seq (scRNA-seq) (Buchon et al., 2013; Hung et al., 2020; Marianes and Spradling, 2013). However, although larval midgut was reported to contain distinct domains in an *in vivo* reporter screen decades ago (Murakami R, Matsumoto A, Yamaoka I, Tanimura T.), its spatial gene expression profile was largely unknown. Here, our Stereo-seq data validated that the embryonic and larval midgut exhibits spatially distinct subregions, indicated by different marker gene expression patterns. Specifically, genes encoding digestive enzymes (including trypsin, amylase, and chymotrypsin) that were regional markers in adult midgut, were also expressed in specific subregions along the embryonic and larval midgut **(Fig.2)**. For instance, in a representative L3 section, the expression of serine protease genes *αTry, βTry, γTry, Jon25Bi* and *Jon25Bii* were localized to a midgut subregion different from that of serine protease gene *CG11911* **(Fig.2E)**. In a representative L2 section, the expression of serine protease genes *βTry, γTry* and *Jon25Bi* were localized to a midgut subregion different from that of starch digestive enzyme genes *Amy-p* and *Amy-d*, and serine protease gene *CG11911* **(Fig.2D)**. Similarly, in a representative L1 section, the set of digestive enzyme genes were also expressed in spatially distinct midgut clusters **(Fig.2C)**. Furthermore, as early as in E14-16 and E16-18 late-stage embryos, spatial expression patterns of digestive enzyme genes were already established **(Fig.2A-B)**, suggesting the maturation of presumptive midgut during embryonic organogenesis. To further reveal the regional diversity of embryonic midgut, we performed Gene Ontology (GO) enrichment analysis on marker genes of each midgut subcluster in E14-16. Marker genes of each subcluster were enriched for distinct functions **(Fig.S5B)**, suggesting that functional regionalization in the midgut already occurs as early as stage 16 of embryogenesis.

**Figure 2.**
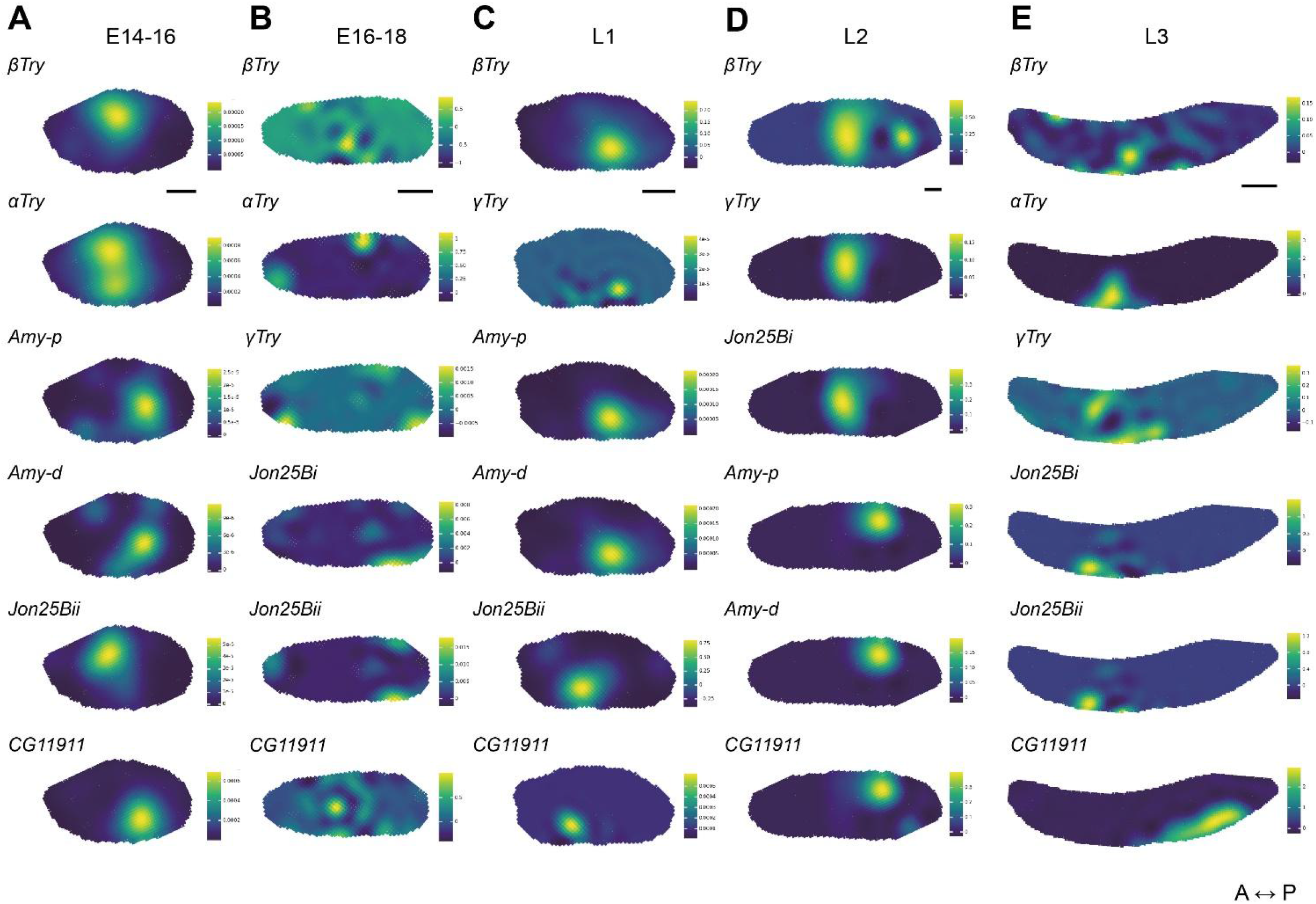
Spatial heterogeneity of digestive enzyme gene expression in midgut across *Drosophila* development. Spatial expression patterns of genes encoding digestive enzymes (trypsin, amylase, and chymotrypsin) in clusters annotated as midgut in representative single sections of **(A)** E14-16, **(B)** E16-18, **(C)** L1, **(D)** L2, and **(E)** L3. All samples are presented in near dorsal/ventral view. Scale bars = 100 μm for E14-16, E16-18, L1 and L2, 500 μm for L3.

Together, our Stereo-seq data uncovered the dynamic spatiotemporal characteristics of functional genes along the digestive tract across *Drosophila* developmental and provided clues for midgut maturation timepoint during embryogenesis.

### Spatiotemporal gene expression and cell state dynamics in reproductive and neural systems

Upon further examination of spatiotemporal features of embryonic and larval tissue transcriptomes, clusters representing male reproductive organs and central nervous system (CNS) stood out as two other regions of high cell type complexity and mRNA abundance. Thus, we extended subclustering and spatial gene expression analysis to these two tissues.

Similar to midgut clusters, we performed further subclustering in clusters representing L3 testis and adjacent clusters expression male germline related genes, and were able to identify clusters corresponding to spermatocytes at different stages of development, distinguished by their marker gene expression pattern (Zhao et al., 2010). To identify the subcluster type, we compared marker gene expression in the clusters representing male reproductive organs with previously published single-cell transcriptomes of L3 gonads subpopulations (Mahadevaraju et al., 2021). Unsupervised clustering enabled identification of spermatogonia and early, mid, and late primary spermatocyte **(Fig.3A)** with distinct marker genes **(Fig.S6)**. Notably, cell type specific marker genes showed clear expression patterns within testis **(Fig.3B)**, reflecting the cell states of spermatocytes at different developmental stages.

**Figure 3.**
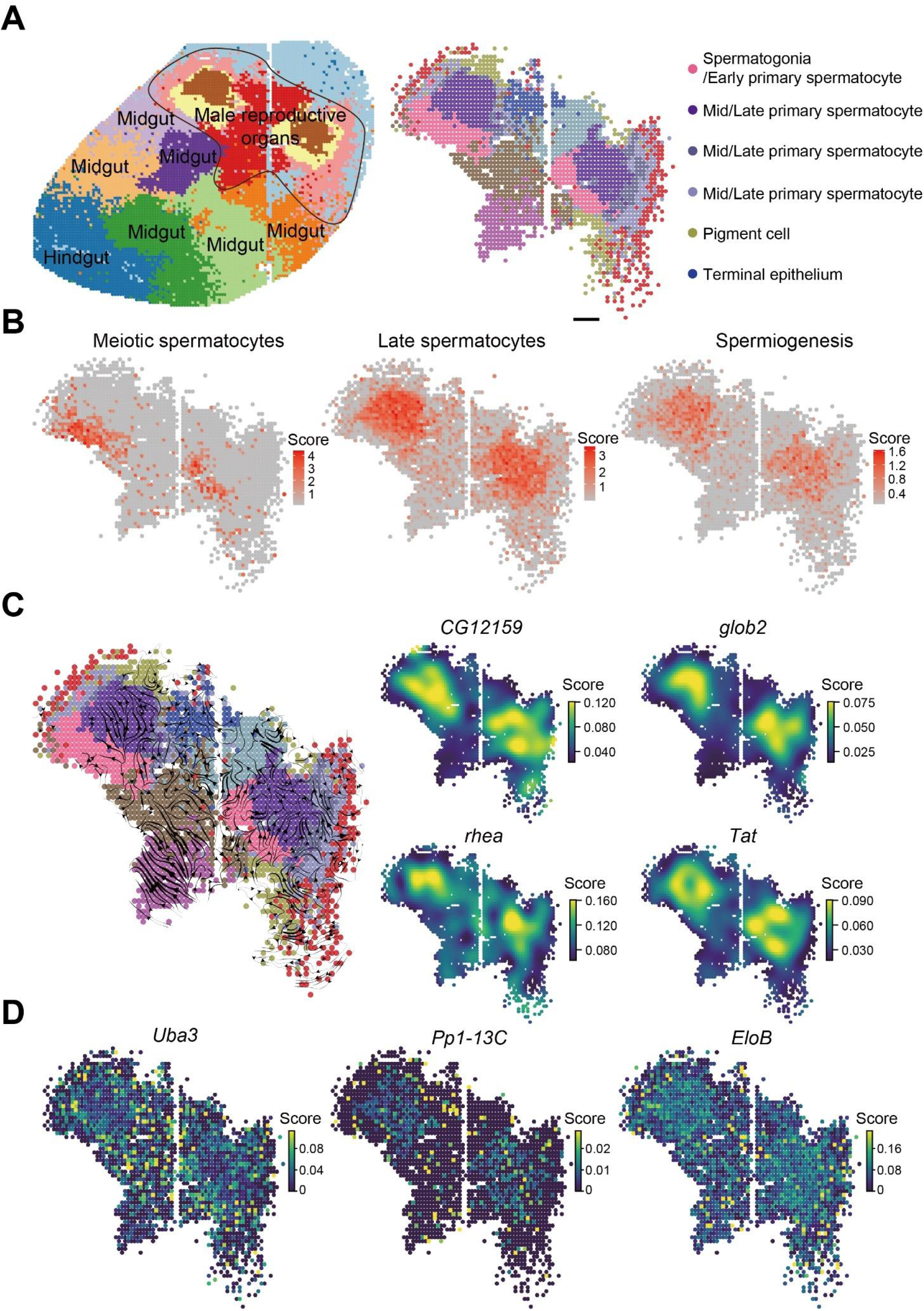
Cell type subclustering and cell state change tracking of *Drosophila* 3rd instar larval male reproductive organs. **(A)** Subclustering results of clusters annotated as testis and adjacent clusters expression male germline related genes in a representative L3 single slice. Scale bar = 100 μm. **(B)** Spatial expression patterns of marker gene sets representing cells at different stages of spermatogenesis (meiotic spermatocytes: *Rbp4, Taf12, blanks*; late spermatocytes: *tHMG1, CG2955, CG13476, CG4375, Pen, SAK, lobo, CG16719*; spermiogenesis: *p-cup, Pif2, whip, CG4073, CG31128, boly, CG14294, hale, Glut3, c-cup, CG12126, CG15059, CG6333, CG31797, CG30099, CG31404, CG13337, CG31815, CG10512, CG14546, CG8851, CG30110*) (Zhao et al., 2010). **(C)** RNA velocity analysis of clusters representing male reproductive organs and expression patterns of representative genes with high velocity. **(D)** Spatial expression patterns of genes encoding inhibitors of Hedgehog and JAK-STAT pathways in male reproductive organs.

In light of the effective capture of cell states in different stages of spermatocytes, we employed RNA velocity analysis (La Manno et al., 2018) to further investigate cell state dynamics. To perform RNA velocity analysis, unspliced mRNAs captured alongside with mature mRNAs by oligo dT primers through secondary priming were first identified. Their abundance was then compared with spliced mature mRNAs and their dynamics were used to infer the directions to which transcript splicing and cell states will change. With RNA velocity analysis, we observed the spatial transcriptional dynamics of spermatocyte maturation in testis **(Fig.3C)**. We then profiled the expression patterns of genes whose mRNAs display high velocity (i.e., undergo rapid transcriptional changes). We found several highly dynamic genes expressed in mid or late spermatocytes, such as Globin gene *glob2. glob2* has been reported to display an unexpected high level of expression in testis, possibly to relieve oxidative stress accumulated during spermatogenesis (Gleixner et al., 2012).

We then profiled the spatial expression patterns of genes involved in Hedgehog or JAK-STAT signaling, which regulate stem cells behavior in the *Drosophila* testis (reviewed in Zhang et al., 2013). We found that transcripts encoding some inhibitors of these pathways, including *Uba3* (Du et al., 2011), *Pp1-13C* (Su et al., 2011), and *EloB* (Stec et al., 2013), were mainly enriched in spermatocytes **(Fig.3D)**, possibly to inhibit the activity of stem cell regulatory pathways in the differentiated spermatocytes.

We also inspected the spatial expression patterns of marker genes in cell subtypes in CNS. The spatial expression patterns of neuronal and glial cell marker genes largely overlap, with glial cell markers showing broader expression patterns, consistent with the spatial distribution of their anatomical structures **(Fig.S7)**. Moreover, we looked at the spatial expression of marker genes of peptidergic and dopaminergic neurons in L2 and found that their spatial patterns can be clearly distinguished **(Fig.S7)**.

To summarize, the high sensitivity of Stereo-seq enabled delicate subclustering of complex tissues to identify cell subtypes in their morphological background. Spatial RNA velocity analysis could provide evidence for activation timing of genes with spatiotemporally restricted regulatory functions.

### Spatial gene regulatory networks during *Drosophila* development

One major driving force of cellular heterogeneity during development is the changes in gene regulatory networks (GRNs), mediated by the interactions of transcription factors (TFs), co-factors and their downstream target genes. Spatial transcriptomic data from Stereo-seq provides a unique opportunity to analyze GRNs in their morphological contexts, allowing GRN identification and analysis in a spatially resolved tissue-specific manner.

To profile the spatial GRNs across *Drosophila* development, we employed single-cell regulatory network inference and clustering (SCENIC) protocol, which were developed to reconstruct GRNs and identify cell states from scRNA-seq data (Aibar et al., 2017). To find regulons of a TF of interest, SCENIC first identifies genes co-expressed with the TF, then enriches for direct-binding downstream target genes with *cis*-regulatory motif analysis. Finally, SCENIC scores the activity of regulons in each cell to identify the cells with high sub-network activity.

SCENIC analysis of Stereo-seq spatial transcriptomic data revealed GRNs with spatial distribution patterns. Across embryonic and larval samples, we identified multiple GRNs mediated by previously reported TFs. Some of the GRNs are tissue-specific, and their corresponding TFs are known to play regulatory roles in the tissues where the GRNs are located, including *GATAe* in midgut (Okumura et al., 2005, 2016), *Pdp1* in muscle (Lin et al., 1997), and *l(3)neo38* (Schneiderman et al., 2010), *kn* (Seecoomar et al., 2000) and *jim* (Iyer et al., 2013) in CNS **(Fig.4)**. Other GRNs are not restricted to certain tissues, but display similar patterns over development, such as *ham, srp, grh* and *Hr78* **(Fig.4)**. The spatial GRN patterns of all these TFs correlate with those of late-stage embryo *in situ* hybridization results **(Fig.4)**, suggesting that SCENIC analysis of Stereo-seq data effectively recapitulated the spatial GRN patterns of these TFs. Among the identified TFs, *ham* was known for its regulatory role in central and peripheral nervous system, and external sensory neurons (Moore et al., 2002, 2004). Some of the examined slices in which *ham* GRNs were identified did not contain CNS, but *ham* GRNs consistently mapped to midgut in late-stage embryos and larvae **(Fig.4)**. It is known that in larvae, *ham* also shows expression enrichment in midgut in addition to CNS (Leader et al., 2018). This suggest that *ham* may play additional regulatory roles in the digestive system.

**Figure 4.**
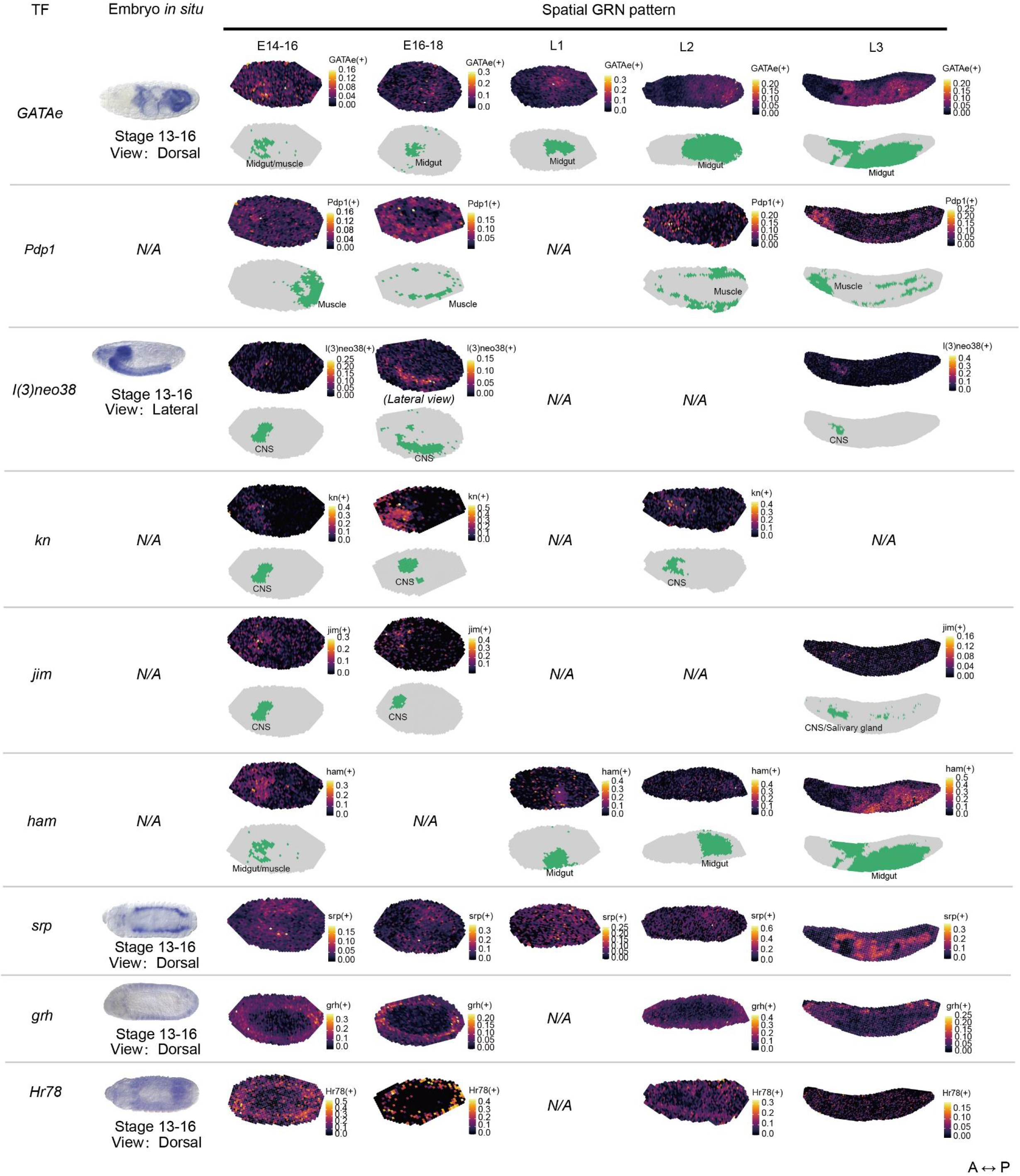
Spatial gene regulatory networks identified in Stereo-seq data. Spatial GRN patterns identified by SCENIC. For each row with two sub-rows, the upper sub-row shows GRNs mapped to spatial transcriptomic profiles generated by Stereo-seq, and the lower sub-row shows the positions and annotations of the clusters co-colocalized with the GRN in the same sample. The left-most column shows *in situ* hybridization results of the TFs from BDGP database (if available). Stages and viewpoints of *in situ* hybridization results are labeled but do not completely match those of our embryo samples. Unless otherwise labeled, Stereo-seq samples are presented in near dorsal/ventral view. N/A indicates that spatial GRN patterns of the TF were not identified in any examined samples of the indicated stage, likely due to incomplete spatial transcriptome capture resulting from cryosection position and orientation. A-P: anterior-posterior.

Thus, SCENIC analysis of Stereo-seq data from developing *Drosophila* identifies both known and potential GRNs, indicating that Stereo-seq can effectively capture transcription regulation networks in the spatial contexts of TFs.

### 3D reconstruction of spatial transcriptome of *Drosophila* late-stage embryos

With the efficient and faithful capture of 2D spatial transcriptome of *Drosophila* embryos by Stereo-seq, we wondered if it is possible to 3D reconstruct the spatial transcriptome of the whole embryo. As a proof of principle, we collected all the 7-μm thick cryosection slices (14 slices obtained) of one entire E16-18 embryo sectioned along the left-right axis (so that each section is a sagittal plane) and performed Stereo-seq with them. We then combined all 2D expression matrices of these samples, performed clustering and annotation individually in each slice **(Fig.S8)**. We developed a pipeline for 3D reconstruction (see **Methods**). Briefly, we extracted 2D sample regions from visualized Stereo-seq expression matrices and aligned them based on shape and transcriptome similarities, so that each bin was assigned an x-y-z 3D coordinate. We then minimized batch effects of each section and merged identical clusters in adjacent slices. The contour and position of 3D clusters resembled the anatomical structures of embryos **(Fig.5A and Supplemental Data 1)**. The marker genes of each tissue displayed expected spatial distribution around their corresponding tissues **(Fig.5B)**. In the 3D reconstructed model, various tissues, represented by clusters aligned across sections, display intra-tissue structural continuity as well as inter-tissue gene expression heterogeneity **(Fig.5C and Supplemental Data 2)**. Moreover, the 3D clusters representing complex tissues such as foregut and midgut can also be further divided into detailed structures based on subclustering and marker gene identification **(Fig.5D and Supplemental Data 3)**.

**Figure 5.**
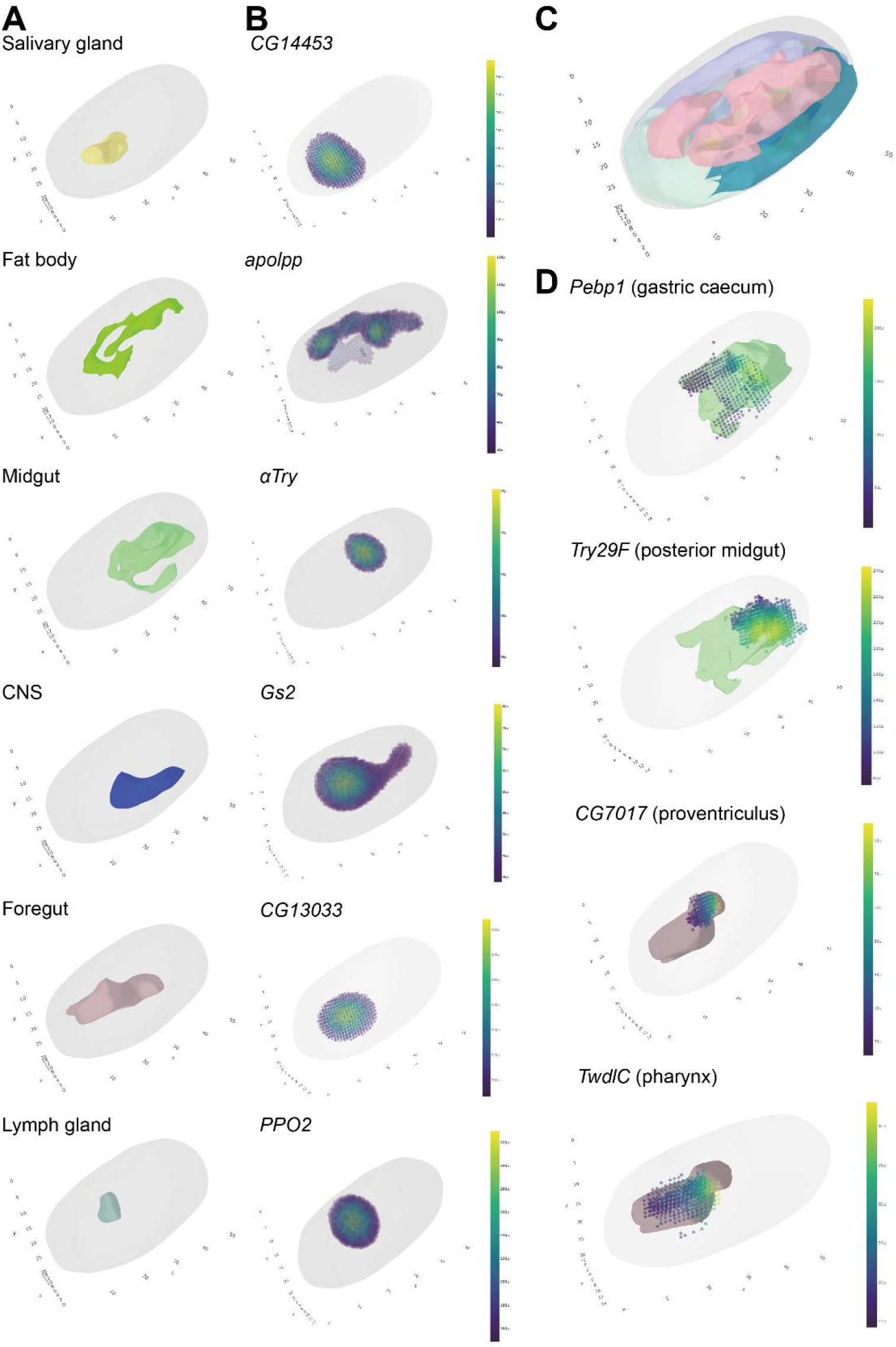
3D reconstruction of spatial transcriptomes of a late-stage *Drosophila* embryo. **(A)** 3D reconstruction, clustering, and annotation of Stereo-seq generated spatial transcriptome of an entire E16-18 embryo. Individual 3D clusters representing specific tissues are shown. **(B)** 3D spatial distribution of representative marker genes of tissues presented in **(A). (C)** The integrated 3D model of the entire embryo. **(D)** 3D clusters representing foregut and midgut, and representative marker genes of their detailed anatomical structures. For scales, each unit in x axis is equivalent to 7 μm (the thickness of a single cryosection slice), and each unit in y and z axis is equivalent to 9.15 μm (the length of 20 DNBs in bin 20 × 20 analysis).

In brief, Stereo-seq is capable of efficient recapitulation of the 3D spatial transcriptome of *Drosophila* embryos. With future optimization of Stereo-seq, we aim to establish an organism-wide, high-resolution, and high-sensitivity 3D spatiotemporal transcriptomic atlas of all stages of *Drosophila* embryos and larvae. We anticipate that with modification, the 3D transcriptome reconstruction strategies we developed here can be applied to additional *Drosophila* adult tissues to resolve their spatial transcriptomes as well.

## DISCUSSION

Spatial transcriptomics techniques greatly expanded our knowledge of gene expression within topographical contexts. In *Drosophila*, existing databases of spatial gene expression patterns were mostly generated from results of *in situ* hybridization analyses. However, *in situ* hybridization techniques are highly limited by probe design strategy and cannot be easily multiplexed. Thus, the gene expression patterns covered by *in situ* hybridization databases are biased. Consequently, previous spatial mapping of *Drosophila* transcriptome and regulatory network is largely based on *in situ* hybridization data and/or computation, which lack the capability for unbiased and global gene expression profiling within intact spatial background.

We present spatiotemporal transcriptomic maps in *Drosophila* from late-stage embryos to 3^rd^ instar larvae based on our Stereo-seq platform. With its unique combination of DNA nanoball patterned arrays and unbiased *in situ* mRNA capture, Stereo-seq provides significantly enhanced resolution and sensitivity compared to other spatial transcriptomics techniques, allowing its application to small-sized tissue sections such as *Drosophila* embryos to simultaneously resolve their spatial transcriptome and morphology. The highly customizable sizes of Stereo-seq chips enable simultaneous mapping of multiple tissue sections, avoiding batch effects introduced by separate experimental runs.

From embryos to larvae, *Drosophila* tissues undergo fundamental diversification in morphology and functions as they develop, which are dictated by spatial (within or between tissue primordia) and temporal (across developmental time) heterogeneity in gene expression profiles. With spatiotemporally resolved gene expression patterns generated by Stereo-seq, expression profiles of tissues across embryonic and larval stages can be dissected in both dimensions to comprehensively reveal their changes during development. Recently, the rapid development of scRNA-seq techniques and the joint efforts of *Drosophila* research community have discovered many new and unanticipated cell types across various tissues during *Drosophila* development (Li et al., 2021). Our Stereo-seq platform can help quickly establish the positions of these newly discovered cells *in vivo*, without the need for *in situ* hybridization with multiple probes. With its high resolution and sensitivity, Stereo-seq itself may also lead to the discovery of new cell types within the spatial contexts in tissues of interest.

We show that Stereo-seq data from *Drosophila* embryos and larvae effectively reflected their tissue anatomical structure and spatial heterogeneity of gene expression. The developmental transcriptomic maps we generated complements current *in situ* hybridization data with quantitative spatial information of both known and previously undetected transcripts. Additionally, the high-resolution and high-sensitivity spatiotemporal transcriptomic data allowed RNA velocity and SCENIC analysis with actual instead of computed spatial background, uncovering the dynamics of known and potential spatially defined tissue-specific gene regulatory networks.

Our results demonstrated the capability of Stereo-seq in resolving the spatial transcriptomes of small-sized samples like *Drosophila* embryos. To our best knowledge, the Stereo-seq data we generated here produces the first actual organism level spatial transcriptomic maps of *Drosophila* late-stage embryos and larvae, which can be combined with scRNA-seq data and provide valuable insights for the systematic study of tissue formation paradigms and regulatory network changes during development. With future optimization of Stereo-seq, we aim to expand the current transcriptomic atlas to generate complete organism-wide 3D spatial transcriptomic profiles covering the entire lifespan of developing *Drosophila*. Application of Stereo-seq to additional genotypes and stages of *Drosophila* will establish an encyclopedic spatial transcriptomic database that will be of great interest to the *Drosophila* research community.

## ACKNOWLEDGEMENTS

This work was supported by the Shenzhen Key Laboratory of Gene Regulation and Systems Biology (Grant No. ZDSYS20200811144002008, China) (to W.C. and Y.H.), the Shenzhen Science and Technology Program (Grant No. KQTD20180411143432337, China) (to W.C. and Y.H.), Shenzhen Key Laboratory of Single-Cell Omics (Grant No. ZDSYS20190902093613831, China) (to L.L.) and Guangdong Provincial Key Laboratory of Genome Read and Write (Grant No. 2017B030301011, China) (to H.L.). This work was also supported by China National GeneBank (CNGB). We thank Dr. Mariana Wolfner for helpful comments on the manuscript. Fig.1A was created with Biorender.com.

## AUTHOR CONTRIBUTIONS

M.W., Q.H. and X.W. conceived the idea; Y.L., X.X., W.C., Y.H. and L.L. supervised the work; M.W., Q.H. and Q.L. designed the experiment; M.W., Q.H., Q.L., Y.W. and Y.A. performed the experiments; T.L., Y.W., Z.T., R.X., K.H. and X.W. processed and analyzed the data; M.C., J.X. and A.C. helped with Stereo-seq library preparation; H.L., W.L. and S.Z. helped with sequencing; Q.H. and Q.L. wrote the manuscript; all authors read and edited the manuscript.

## DECLARATION OF INTERESTS

The chip, procedure, and application of Stereo-seq are covered in pending patents. Employees of BGI have stock holdings in BGI.

## RESOURCE AVAILABILITY

### Lead contact

Further information and requests for the resources and reagents may be directed to the corresponding author Longqi Liu.

### Material availability

All materials used for Stereo-seq are commercially available.

### Data and code availability

All raw data generated by Stereo-seq and custom codes using open-source software to support this study are provided in supplementary materials (**Supplemental Data 4**) or deposited to CNGB Nucleotide Sequence Archive database with accession code: CNP0002189 (https://db.cngb.org/search/project/CNP0002189). All data were analyzed with standard programs and packages, as detailed in **Methods**.

## METHODS

### Fly strain maintenance

All embryo and larva samples in this study were from *Drosophila* strain *w1118* (Tsinghua Fly Center). Flies were maintained on cornmeal-sucrose-agar media (Hopebio, HB8590) in a 25 °C incubator (Laifu, PGX-280A-3H) on a 12 h/12 h light/dark cycle.

### Sample preparation

Embryos were collected at two-hour intervals on a grape juice plate [2.15% w/v agar (Vetec, V900500), 49% v/v grape juice, 0.2% v/v propionic acid (Aladdin, P110444), 0.02% phosphate acid (LingFeng, Shanghai)] from a population cage and aged to desired stages. Larvae of desired stages were isolated from the same population cage. Samples were incubated in phosphate buffer saline (PBS) containing 0.5 mg/mL bromophenol blue (Macklin, B802656) for 10 min for staining and better visualization during cryosection. Samples were then rinsed in PBS to remove excessive dye before embedding. To compare the efficiency of mRNA capture and 3D reconstruction, we attempted to cut the samples in two directions (along the left-right axis for sagittal section, or along the dorsal-ventral axis for transverse section) during cryosection. The exact orientations of each sample used in this study were indicated in the figure legends. Samples were oriented so that they were most likely to be sectioned in the desired axis. Orientated samples were immobilized with double-sided tapes to prevent disturbance from flowing embedding media. 6 samples of the same stage (2 samples for L2 and L3) were embedded and sectioned together. Sample were embedded with Tissue-Teck OCT (Sakura, 4583) and transferred to a -80 °C freezer for storage until used. Cryosection was performed with indicated slice thickness in a Leica CM1950 cryostat. Sample sections were applied to Stereo-seq chips immediately after cryosection.

### Stereo-seq

Stereo-seq library preparation and sequencing were performed as previously described (Chen et al., 2021). Briefly, embryo and larva sections on Stereo-seq chips were fixed in pre-chilled methanol at -20 °C for 40 min. After removal of methanol, sections were permeabilized on chip with 100 μl 0.1% pepsin (Sigma, P7000) in 0.01 M HCl at 37 °C for 5 min. Permeabilization solution was then removed and sections were washed with 100 μl 0.1× saline-sodium citrate (SSC, Thermo, AM9770) buffer supplemented with 0.05 U/μl RNAase Inhibitor (MGI). mRNAs captured by DNBs on the chip were reverse transcribed with SuperScript II reverse transcription (RT) mix (Invitrogen, 18064-014, 10 U/μL reverse transcriptase, 1 mM dNTPs, 1 M betaine solution PCR reagent, 7.5 mM MgCl_2_, 5 mM DTT, 2 U/μL RNase inhibitor, 2.5 μM Stereo-seq template switch oligo and 1× First-Strand buffer) at 42 °C for 1 h. RT mix was then removed. The chip was washed with 0.1× SSC and incubated in tissue removal buffer (10 mM Tris-HCl, 25 mM EDTA, 100 mM NaCl, 0.5% SDS) at 37 °C for 30 min. Tissue removal buffer was removed and the chip was washed twice with 0.1× SSC. cDNAs on the chip were amplified with KAPA HiFi Hotstart Ready Mix (Roche, KK2602) with 0.8 μM cDNA-PCR primer. Sequencing libraries were prepared with PCR products undergoing the following steps: fragmentation (in-house Tn5 transposase), amplification (KAPA HiFi Hotstart Ready Mix), and purification (Vazyme, N411-03). Final libraries were sequenced on a MGI DNBSEQ-Tx sequencer.

### Stereo-seq data analysis

#### Raw data processing

Stereo-seq raw data processing and unsupervised clustering were performed as previously described (Chen et al., 2021). Briefly, CID sequences were first mapped to the designed coordinates on chip with 1 base mismatch tolerance. MID sequences with quality score lower than 10 were filtered out. cDNA sequences were aligned to the reference genome (*Dm3*) by STAR (Dobin et al., 2013). Expression profile matrices with CID were generated based on the information above.

#### Unsupervised clustering

The expression profile matrices of sections from embryos and L1/L2 were divided into bins with 20 × 20 DNBs, while those from L3 were divided into bins with 50 × 50 DNBs. After obtaining the expression profile matrices through the in-house processing software, we used *Seurat* package to perform data normalization and unsupervised clustering (Stuart et al., 2019). Briefly, *SCTransform* function was first applied to normalize and identify highly variable genes. After dimensionality reduction with *runPCA* function, *runUMAP* function was used to perform a two-dimensional projection on the data with the following parameter: *dims = 6*. The identification of each cluster was then performed with *FindNeighbors, FindClusters*, and *FindAllMarkers* functions in *Seurat* with the following parameters: *dims = 6, verbose = FALSE, resolution = 0*.*6, only*.*pos = TRUE, min*.*pct = 0*.*25, logfc*.*threshold = 0*.*25*.

#### Cluster annotation

For the list of marker genes of each cluster in the unsupervised clustering results, we extracted the top 30 genes with the most significant *p* values. To infer specific tissue types of each cluster, we queried these 50 genes in the publicly available databases, including BDGP, FlyBase, FlyAtlas1, FlyAtlas2 and FlyMine, as well as other published results. We assigned the tissue type(s) that match the most marker genes of the cluster.

### RNA velocity analysis

Spliced and unspliced RNA were counted for all detected genes (∼400 genes per bin in the male reproductive organ clusters) by every bin 50 × 50 using Velocyto command line interface (La Manno et al., 2018), with about 1.3% of genes per bin were detected as unspliced type. The generated splice and unspliced expression matrices were then used to estimate the differentiation dynamics of cell lineages with dynamo software (Qiu et al., 2021), the transitioning between gene expression states was modeled and inferred, with lineage relationship of early primary spermatocyte to mid primary spermatocyte set as confident cell velocities, to determine the expression dynamics of testis cell. Genes with high correlation with the transition were also determined in the same process.

### SCENIC analysis

*pySCENIC* pipeline (Van de Sande et al., 2020) was used to predict activity scores of TFs on each section. *pySCENIC* uses TFs’ motif and downstream gene expression to predict TFs’ activity scores at single cell level. *pySCENIC* pipeline was first used to generate AUC matrices of TFs, with rows representing bins and columns representing TFs. Values in each cell represent TFs’ activity scores. AUC values in each section were then visualized with *ggplot2* (Wickham, 2011).

### Gene Ontology enrichment analysis

R package *clusterProfiler* (Yu et al., 2012) was used to identify enriched Gene Ontology terms. Marker gene lists from each cluster were inputted with default parameters.

### Imputation of spatial gene expression

For better visualization of spatial gene expression patterns, *SparseVFC* function of dynamo software was used to impute gene expression by 2D or 3D coordinates to render spatial patterns of gene expression. This method was used in **Fig. 2, Fig.3C** and **Fig.S7**.

### 3D Reconstruction of embryo transcriptome

#### Stereo-seq data preprocessing

2D expression matrices of all 14 sections of an entire E16-18 embryo were subjected to preprocessing. Specifically, for each section, Python package *scanpy* (Wolf et al., 2018) was used to normalize expression values for total UMI counts per bin 20 × 20. *calculate_qc_metrics* function was used for data quality control, with the following non-default parameters: *percent_top = None, log1p = False, inplace = True*. Highly variable genes were identified by function *highly_variable_genes* with parameters: *flavor = seurat, n_top_genes = 2000, inplace=True*. Other functions in preprocessing used standard methods and default processes of *scanpy*, including total-count normalization and principal component analysis.

#### Unsupervised clustering and cluster annotation

For each section, PCA algorithm was used for data dimensionality reduction with parameter *n_comps = 20*. Python package *squidpy* (Palla et al., 2021) algorithm was used for unsupervised clustering. Specifically, after using *neighbors* function in *scanpy* to find the neighbors of each bin 20 × 20 based on gene expression information with parameter *n_neighbors = 30, spatial_neighbors* function in *squidpy* was used to find the spatial neighbors of each spot with parameter *n_neigh = 6*. The neighbor information obtained through the two algorithms were then merged. Based on the merged neighbor information, *leiden* algorithm in *scanpy* was used for unsupervised clustering. After individual testing, optimal resolution was determined for each section. Clusters in the optimal resolution of each section were annotated as described above.

#### Section alignment and 3D modeling

For two adjacent sections, PASTE (Zeira et al., 2021) algorithm was used to align them along the z axis based on both gene expression similarities and spatial coordinates. For all the 2D expression matrices of 14 sections, *pairwise_align* function in PASTE was run sequentially along the z axis, and each bin was assigned an x-y-z 3D coordinate. After that, on 2D graphics of all sections, bins with the same annotation were assigned the same color code. The aligned 3D coordinates and color codes representing different tissues were integrated into 3D graphics in tiff format with *skimage* (Van der Walt et al., 2014) algorithm. *3D Slicer* (Fedorov et al., 2012) software was used to transform tiff data into a smooth 3D model of the embryo. Major smoothing and 3D modeling methods used were *Margin, Closing, Opening, Median, Gaussian*, and *Joint smoothing*. Spatial distribution of 3D clusters and gene expression patterns were visualized with *scanpy* and *plotly* (Sievert, 2020). Python tools *matplotlib* (Hunter, 2007) and *plotnine* (Kibirige, 2017) were used to draw 2D graphics, and *plotly* was used to display 3D graphics and models.

## FIGURE LEGENDS

**Figure S1.**
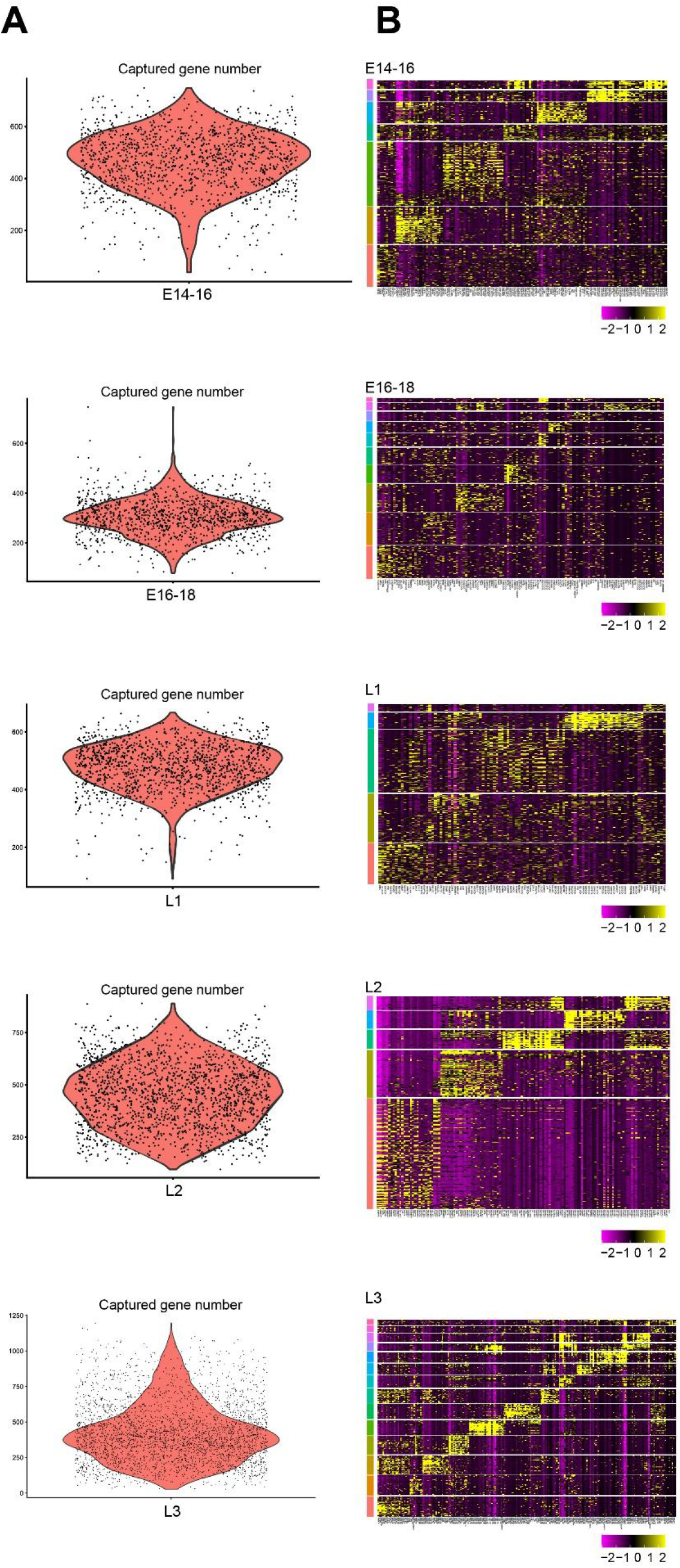
Quality control of Stereo-seq data from samples of each examined stages. **(A)** Numbers of genes captured by Stereo-seq in representative samples, each dot represents one bin (bin 20 × 20 for E14-16, E16-18, L1 and L2, bin 50 × 50 for L3). **(B)** Heatmap of expression levels of top marker genes of each cluster after unsupervised clustering in the same samples as **(A)**.

**Figure S2.**
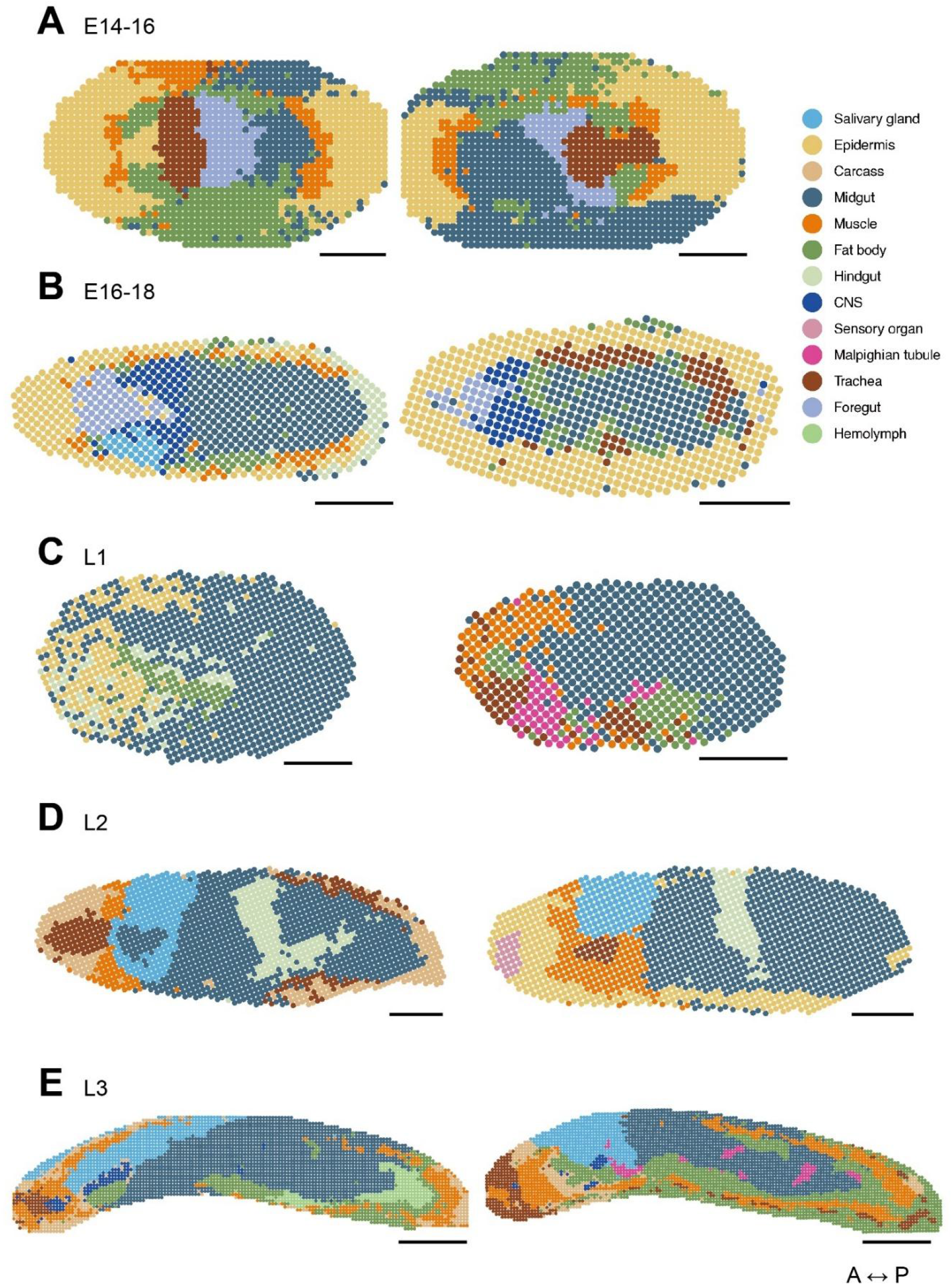
Biological replicates of spatial transcriptomes of *Drosophila* embryos and larvae generated by Stereo-seq. **(A-E)** Unsupervised clustering and annotation of single sections of **(A)** E14-16, **(B)** E16-18, **(C)** L1, **(D)** L2, and **(E)** L3 samples. Clusters with the same annotation are merged and assigned the same color code. Scale bars = 100 μm for E14-16, E16-18, L1 and L2, and 500 μm for L3. A-P: anterior-posterior.

**Figure S3.**
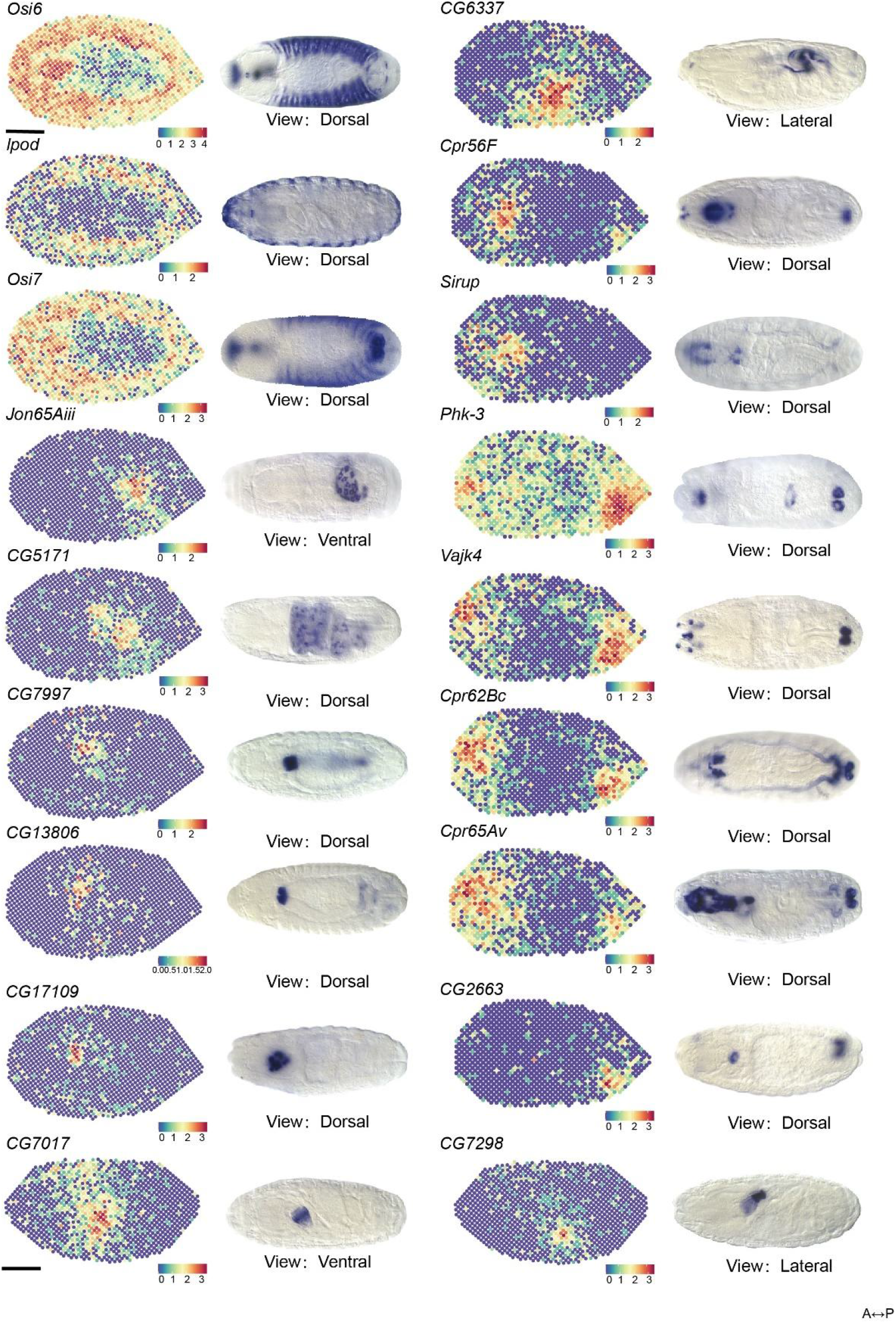
Comparisons of spatial gene expression patterns between Stereo-seq and *in situ* hybridization. For each gene panel, left shows spatial expression patterns generated from Stereo-seq data of representative single sections of E14-16 or E16-18 embryo samples; right shows *in situ* hybridization results of embryos at similar stages from BDGP database. All Stereo-seq samples are in near dorsal/ventral view. Viewpoints of each *in situ* hybridization image are labeled under each panel. Scale bars = 100 μm. A-P: anterior-posterior.

**Figure S4.**
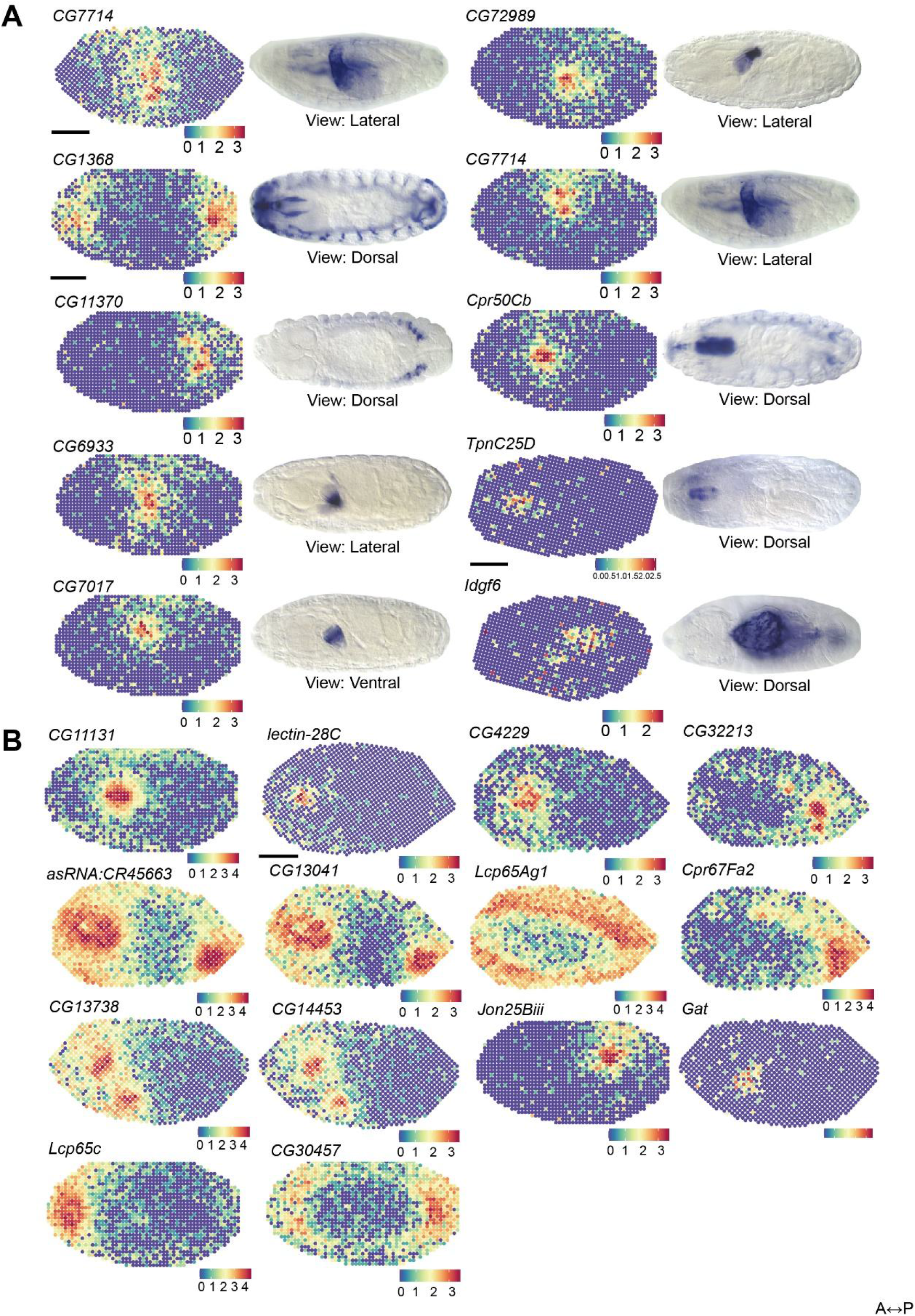
Comparisons of spatial gene expression patterns between Stereo-seq and *in situ* hybridization. **(A)** Continued from **Figure S3. (B)** Spatial expression pattern of genes with no available clear *in situ* hybridization patterns from BDGP or Fly-FISH. All Stereo-seq samples are in near-dorsal/ventral view. Viewpoints of each *in situ* hybridization image are labeled under each panel. Scale bars = 100 μm. A-P: anterior-posterior.

**Figure S5.**
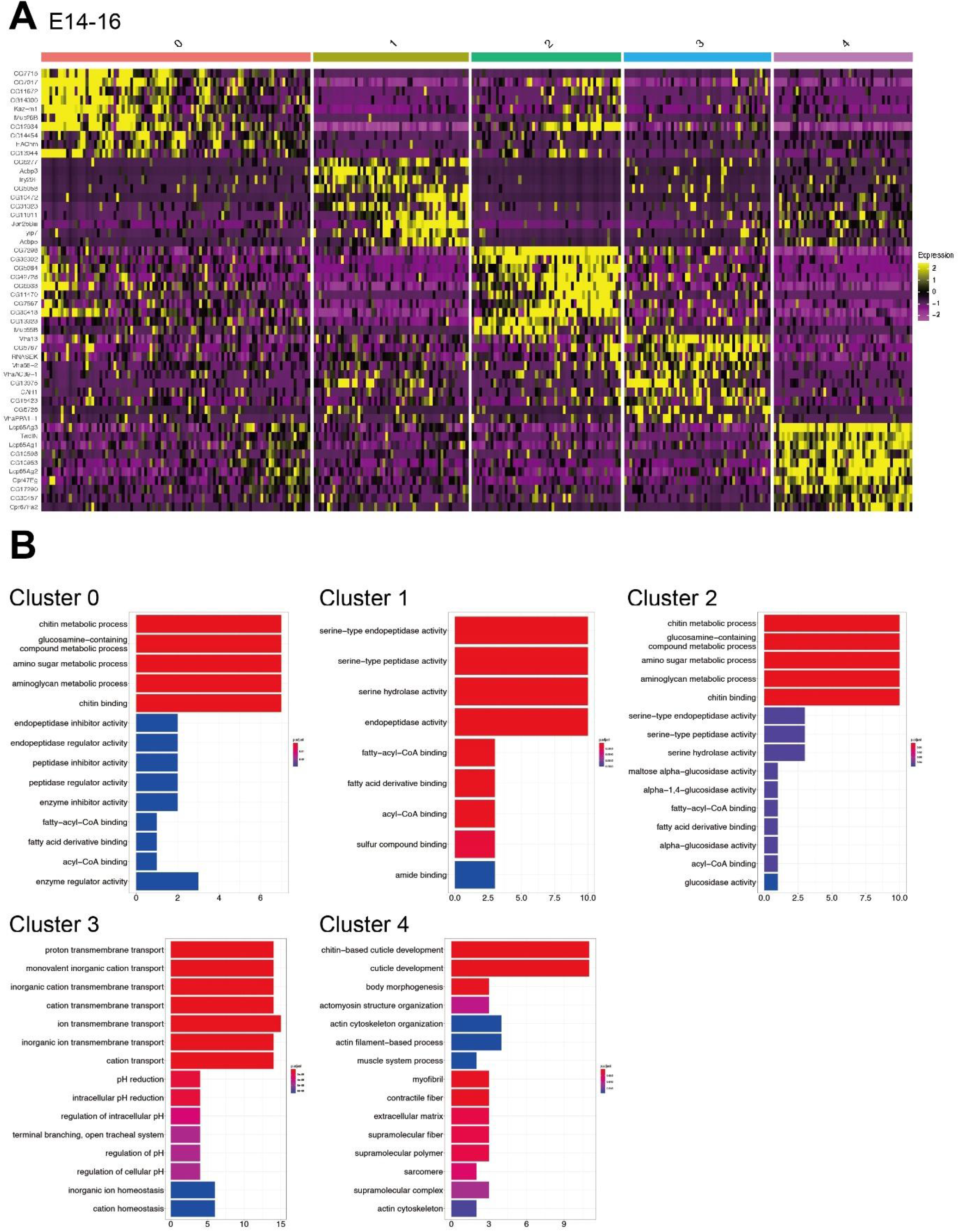
Analysis of marker genes of embryonic and larval midgut subregions. **(A)** Heatmap of the top marker genes of midgut subclusters in the single section from E14-16 in **Figure 2. (B)** GO ontology analysis of marker genes of each subcluster in **(A)**.

**Figure S6.**
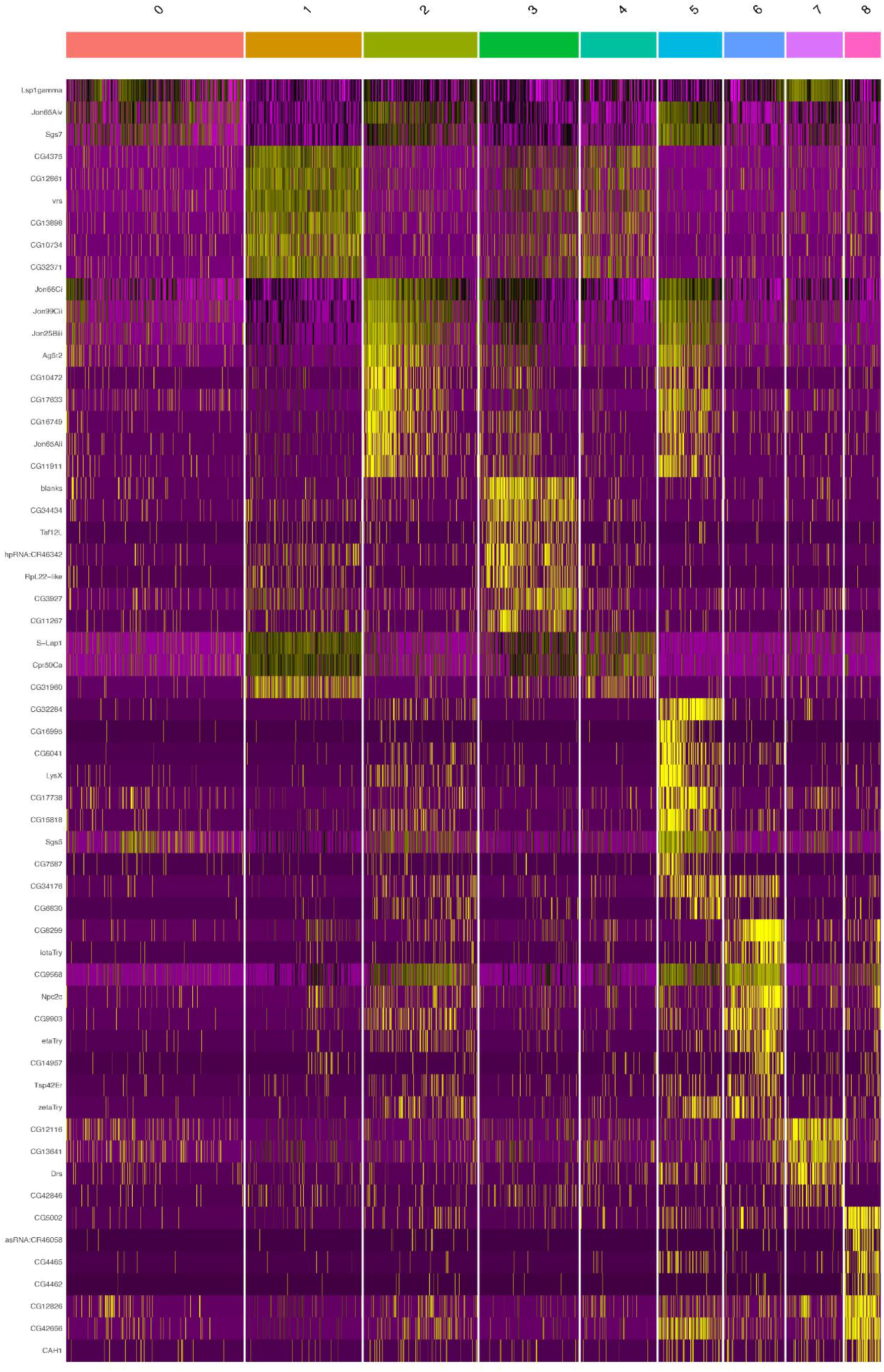
Heatmap of the top marker genes of subclusters of male reproductive organs in the single L3 section in Figure 3.

**Figure S7.**
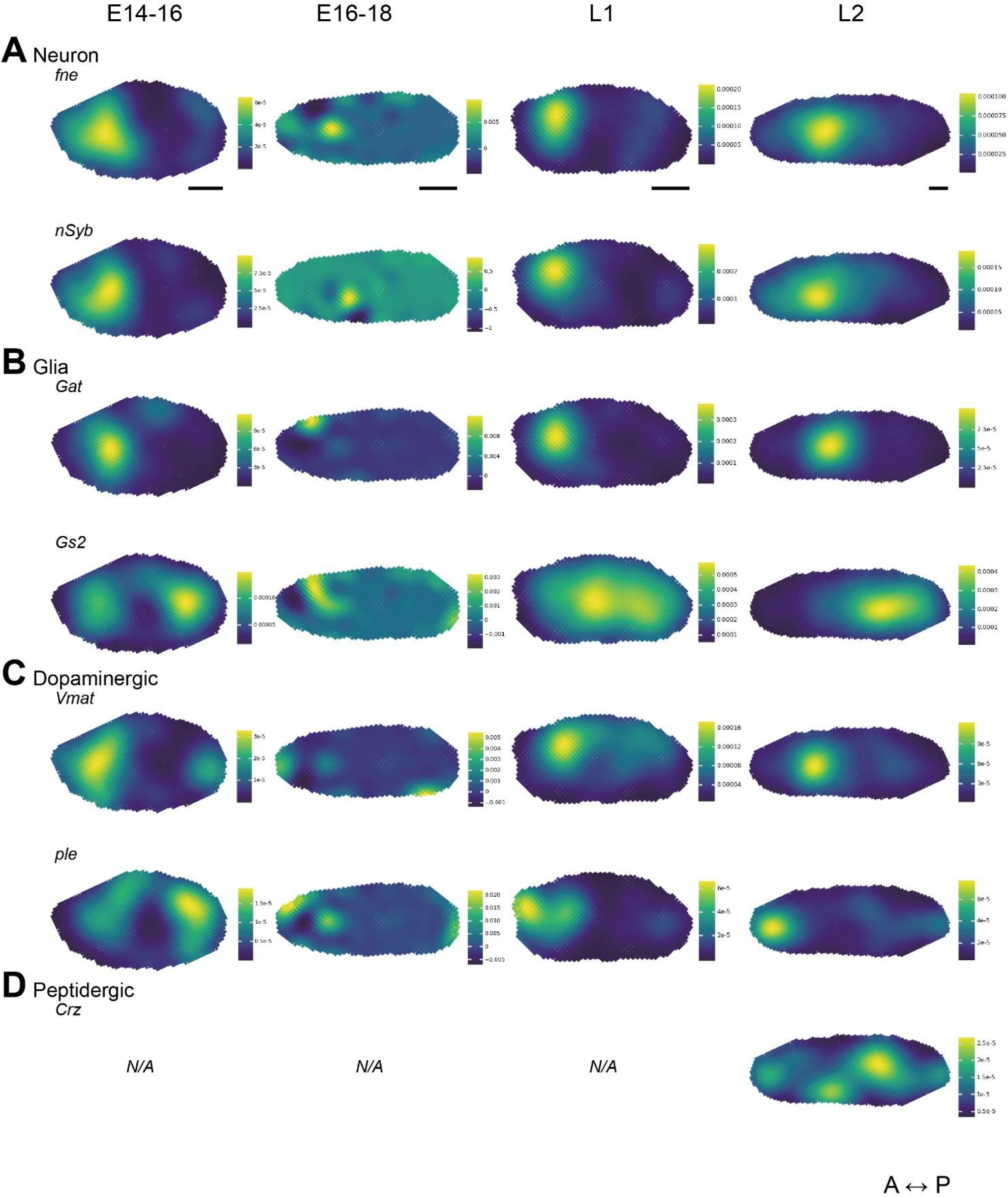
Spatial expression patterns of marker genes of embryonic and larval CNS. **(A-D)** Spatial expression patterns of marker genes of **(A)** neural, **(B)** glial cells, **(C)** peptidergic, and **(D)** dopaminergic neurons in late-stage embryos and larvae. N/A indicates that spatial gene expression patterns were not identified in any examined samples of the indicated stage, likely due to incomplete spatial transcriptome capture resulting from cryosection position and orientation. Scale bars = 100 μm. A-P: anterior-posterior.

**Figure S8.**
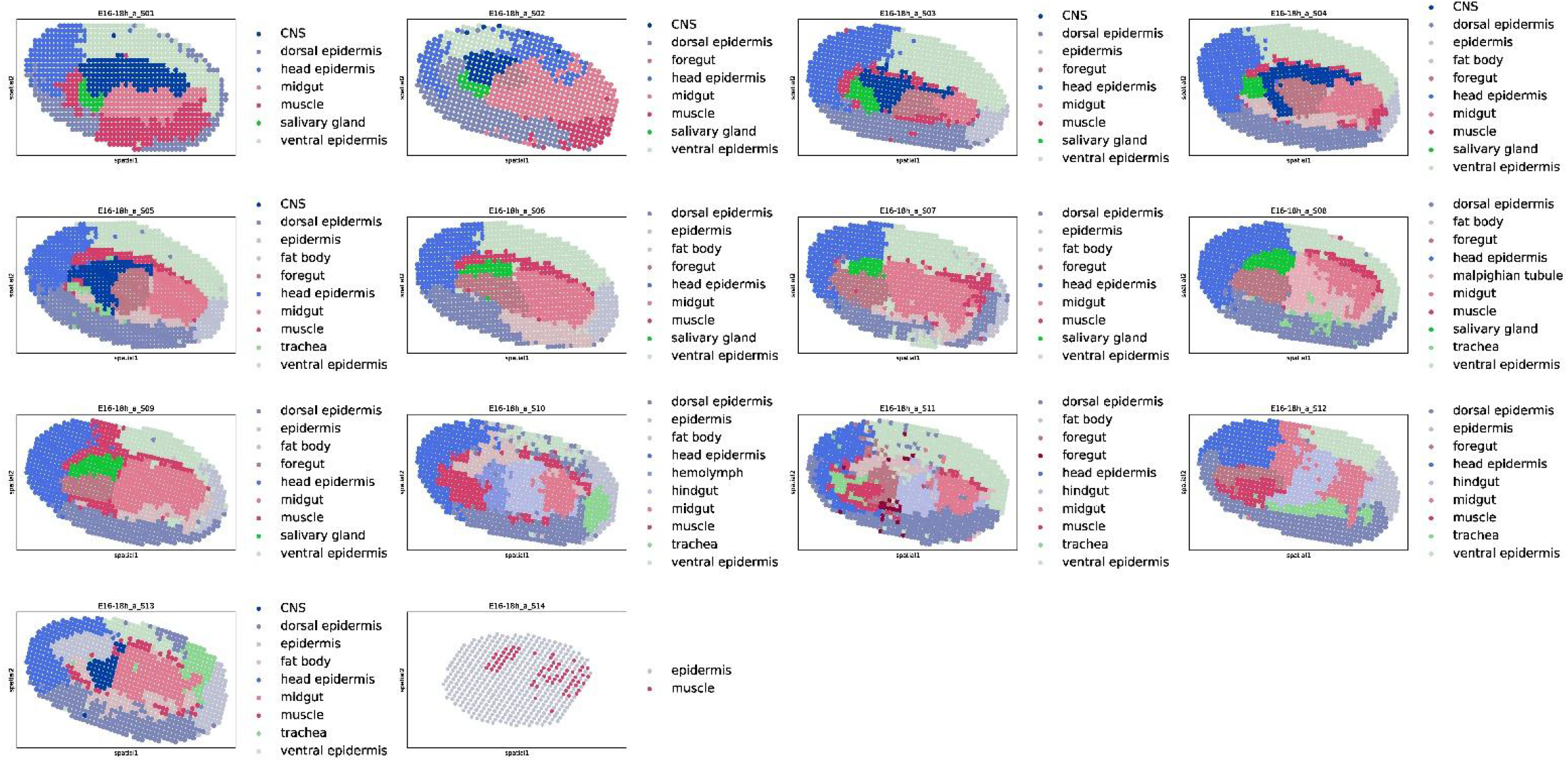
Clustering and annotation of Stereo-seq generated spatial transcriptome of all sections of an E16-18 embryo used for 3D reconstruction in Figure 5.

**Table S1.** Numbers of transcripts and genes captured in samples shown in Figure 1. Numbers are shown in mean ± standard deviation.

| Sample | Bin size (number of DNBs) used for analysis | Number of unique transcripts per bin | Number of unique genes per bin |
| --- | --- | --- | --- |
| E14-16 | 20 × 20 | 1325.2 ± 347.0 | 437.2 ± 81.3 |
| E16-18 | 20 × 20 | 2969.1 ± 913.9 | 425.7 ± 78.9 |
| L1 | 20 × 20 | 1602.1 ± 368.4 | 477.9 ± 82.4 |
| L2 | 20 × 20 | 1928.3 ± 923.7 | 453.9 ± 151.4 |
| L3 | 50 × 50 | 1768.4 ± 1737.7 | 423.8 ± 208.5 |

**Supplemental Data 1** 3D spatial patterns of representative tissues in an entire E16-18 embryo shown in **Figure 5**.

**Supplemental Data 2** 3D spatial transcriptome reconstruction model of an entire E16-18 embryo shown in **Figure 5**.

**Supplemental Data 3** 3D spatial patterns of representative marker genes of detailed anatomical structures in foregut and midgut in an entire E16-18 embryo shown in **Figure 5**.

**Supplemental Data 4** Custom scripts used in this study.

## Notes

### Competing Interest Statement

The authors have declared no competing interest.

